# Impact of high phosphorous and sodium on productivity and stress tolerance of *Arundo donax* plants

**DOI:** 10.1101/477810

**Authors:** Claudia Cocozza, Federico Brilli, Laura Miozzi, Sara Pignattelli, Silvia Rotunno, Cecilia Brunetti, Cristiana Giordano, Susanna Pollastri, Mauro Centritto, Gian Paolo Accotto, Roberto Tognetti, Francesco Loreto

## Abstract

*Arundo donax* L. is an invasive species recently employed for biomass production that emits large amounts of isoprene, a volatile compound having important defensive role. Here, the potential of *A. donax* to grow in degraded soils, characterized by poor fertility, eutrophication and/or salinization, has been evaluated at morphological, biochemical and transcriptional level. Our results highlight sensitivity of *A. donax* to P deficiency. Moreover, we show that *A. donax* response to salt stress (high sodium, Na^+^), which impaired plant performance causing detrimental effects on leaf cells ultrastructure, is characterized by enhanced biosynthesis of antioxidant carotenoids and sucrose. Differently from Na^+^, high phosphorous (P) supply did not hamper photosynthesis although it affected carbon metabolism through reduction of starch content and by lowering isoprene emission. In particular, we revealed on salt-stress leaves that high P enhanced the expression of genes involved in abiotic stress tolerance, but further increased diffusive limitations to photosynthesis and slowed-down sugar turnover without modifying isoprene emission. Therefore, despite limiting productivity, high P improved *A. donax* tolerance to salinity by favouring the accumulation of carbohydrates that protect cells and increase osmotic potential, and by stimulating the synthesis of antioxidants that improves photo-protection and avoids excessive accumulation of reactive oxygen species.

**Highlights:** *Arundo donax* is sensitive to elevated salinity. High phosphorous supply to salt-stressed *A. donax* enhances transcriptomic changesthat induce the onset of physiological mechanisms of stress tolerance but limits productivity.

## Introduction

Phosphorus (P) is an essential element for many key enzymes and intermediates of plants photosynthetic CO_2_ assimilation and sugar biosynthesis (Beck and Ziegler, 1989). Phosphorous regulates energy storage reactions and maintains structural integrity of cellular membranes (Marschner, 1995). Phosphorous concentration is tightly regulated within the cells because changes in its availability can seriously impair plant physiological processes and structure (Shen *et al.*, 2011). On one hand, P deficiency affects the overall plant metabolism (Hernández *et al.*, 2007) reducing growth (Chiera *et al.*, 2002) and hampering the ability to reproduce and adapt to different environments (Wassen *et al.*, 2005). On the other hand, P surplus decreases plant performances by inhibiting the biosynthesis of starch (Fredeen *et al.*, 1989) and other secondary metabolites (e.g. isoprenoids, Fernández-Martínez *et al.*, 2017, Fares *et al*., 2008), and lowers nitrate assimilation in the roots (Rufty *et al.*, 1990).

Intensive exploitation of phosphate rock reserves for fertilization purposes may lead to their depletion by the end of this century (Cordell *et al.*, 2009). However, marginal lands, where high amounts of P are associated with salinity, are not suitable for agriculture. It is well-know that salinity impairs plant performance and productivity (Munns and Tester, 2008). In particular, exposure to high sodium (Na^+^) concentration in soil increases diffusive (Centritto *et al.*, 2003) and biochemical limitations to photosynthetic CO_2_ assimilation (Chaves *et al.*, 2009), decreases water transpiration rates, modifies the biosynthesis of both soluble sugars (Dubey and Singh, 1999) and starch (Parida *et al.*, 2002), and reduces pigments content in leaves (Kalaji *et al.*, 2011). Moreover, excess of Na^+^ impairs root nutrient uptake by altering the trans-membrane transport of ions that leads to loss of turgor of plant cells and to further membrane damage following the formation of reactive oxygen species (ROS) (Sobhanian *et al.*, 2011).

*Arundo donax* L., the giant reed, is a non-food perennial rhizomatous invasive grass species belonging to *Poaceae* family (Pilu *et al.*, 2012). *A. donax* is one of the most efficient C3 plant species, able to colonize a wide spectrum of habitats worldwide, from very wet loam to relatively dry sandy soils (Webster *et al.*, 2016). *A. donax* displays a high photosynthetic rate and a fast-growing habit that make its cultivation suitable for biomass and bioenergy production (Webster *et al.*, 2016; Sánchez *et al.*, 2016). In addition, the tolerance to abiotic stress of *A. donax* has been already demonstrated across a range of stressful conditions, thus allowing the exploitation of degraded and marginal lands with *A. donax* crops (Calheiros *et al.*, 2012; Nackley and Kim, 2015). In fact, *A. donax* is able to maintain a high leaf-level photosynthesis and biomass gain under drought (Haworth *et al.*, 2017b) and salinity (Nackley and Kim, 2015). In particular, efficient stomata regulation in *A. donax* is induced by increase in leaf ABA content in response to drought (Haworth *et al.*, 2018). Moreover, *A. donax* is able to adjust the xylem vessel size to regulate the vulnerability to embolism under water deficit conditions (Haworth *et al.*, 2017c). Recently, it has been shown that symbiosis with arbuscular mycorrhiza increases *A. donax* performance to salinity, through proline accumulation and isoprene formation (Pollastri *et al.*, 2018).

*A. donax* leaves constitutively produce a large amount of isoprene (Velikova *et al.*, 2016), which is known to be involved in mechanisms of protection against abiotic (Vickers *et al.*, 2009) and biotic stresses (Loivamäki *et al.*, 2008). However, there is no clear pattern in isoprene emission in response to abiotic stress in reeds, as isoprene emission increased in *A. donax* following drought (Haworth *et al.*, 2017a), decreased in *Phragmites australis* (the common reed, and a close relative of Arundo) plants exposed to high P concentrations (Fares *et al.*, 2006), and was unaltered in salt-stressed *A. donax* (Pollastri *et al.*, 2018).

In this study, *A. donax* plants grew under controlled laboratory conditions by providing a nutrient solution deprived of P, or enriched with a high concentration of P also in combination with high concentrations of sodium chloride (NaCl). Our investigation aimed at: a) characterizing the response of *A. donax* plants to P availability, both under P-deficiency and supply of high P concentration; b) testing the performance of *A. donax* under multiple (abiotic) stresses, such as a simultaneous excess of P and Na^+^. To this purpose, we used an integrated approach, combining physiological and biochemical measurements with transcriptomic analysis. Leaf and root transcriptomes of *A. donax* have been recently explored only in healthy plants (Sablok *et al.*, 2014) and in plants exposed to drought stress (Fu *et al.*, 2016; Evangelistella *et al.*, 2017). Understanding, at molecular level, the response of *A. donax* to combined P and Na^+^ stress is crucial for implementing adaptation strategies in order to achieve high biomass yield and productivity in marginal areas for agriculture.

## Materials and methods

### Plant material, growth conditions, supply of P and Na^+^

*A. donax* plants were propagated from rhizomes of plants collected in Sesto Fiorentino (Italy). Rhizomes were kept in tap water for one day (d) and then planted in 6 dm^3^ pots containing quartz sand. All potted plants were then grown in a climatic chamber under controlled environmental conditions (temperature ranging between 24°C and 26°C; relative air humidity ranging between 40% and 60%; photosynthetic photon flux density (PPFD) of 700 μmol m^-2^ s^-1^ for 14 h per d), and were regularly watered twice a week with half strength Hoagland solution (Hoagland and Arnon, 1950) for two months before beginning the experiment.

The experiment was performed by applying five different nutrient conditions: 1) Hoagland solution (C); 2) Hoagland solution deprived of phosphorous (−P); Hoagland solution complemented with 8.0 mM KH_2_PO_4_ (+P); Hoagland solution complemented with 200 mM NaCl (+Na); and Hoagland solution complemented with both 200 mM NaCl and 8.0 mM KH_2_PO_4_ (+NaP). All different solutions were supplied twice a week during the whole duration of the experiment (43 days).

### Biometrical traits, leaf determination of C, N, P and Na^+^

Biometrical traits (leaf number and culm length) were measured weekly. The relative water content (RWC) of leaves was determined on the second fully expanded leaf at the end of the treatment. Fresh weight (FW) was immediately determined following leaf collection. The same leaf was then immersed into water for 24 h before measuring the turgid weight (TW). Finally, the leaf was oven dried at 80°C for 48 h before measuring the dry weight (DW). RWC was calculated by using the formula:

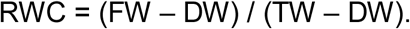

Total C and total N concentrations (%) were quantified at the end of the treatment with a Carlo Erba NA 1500 CNS Analyzer (Milan, Italy) through the chromatographic column by a thermal conductivity detector. Na^+^ and P concentrations were determined at the end of the treatment. Na^+^ concentration was measured by flame atomic absorption spectrometry (Analyst 200, Perkin Elmer), and P concentration was measured by inductively coupled plasma atomic emission spectrometry (ICP-AES; iCAP 6500 Duo; ThermoFisher, Dreieich, Germany), employing appropriate quality standard controls (Sreenivasulu *et al.*, 2017).

### CO_2_/H_2_O gas exchange, fluorescence and isoprene measurements

Gas exchange of CO_2_ and H_2_O and fluorescence measurements were performed at the end of the treatment by using a portable gas exchange system equipped with a fluorometer (Li-Cor 6400, Li-Cor Biosciences Inc., NE, USA). The third (from the shoot apex) fully expanded leaf of *A. donax* was clamped in the 2 cm^2^ Li-Cor cuvette and exposed to a saturating PPFD of 1000 μmol m^-2^ s^-1^, CO_2_ concentration of 400 ppm, temperature of 30°C and relative humidity (RH) ranging between 45 and 50%. Photosynthesis (A), stomatal conductance (g_s_) and internal CO_2_ concentration (Ci) were calculated using the formulations of von Caemmerer and Farqhuar (1981) 10 min after reaching steady-state conditions. The linear electron transport rate (ETR) was calculated from fluorescence measurements of PSII efficiency, according to Genty *et al.* (1989).

Photosynthesis under low O_2_ conditions was measured reducing the air O_2_ concentration from 21% to 2%. We used a nitrogen cylinder connected with a mass flow controller (Rivoira, Italy) that precisely enriched the concentration of N_2_ in the air entering the LI-Cor 6400 from 89 to 99%, while CO_2_ concentration was maintained steady at 400 ppm. The O_2_ inhibition of photosynthesis was calculated from the A values measured at 21% and at 2% of O_2_ (v/v) using the following formula (Zhang *et al.*, 2017):

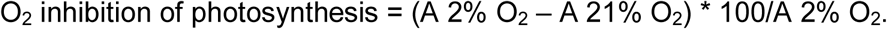

Isoprene emitted by leaves was collected at the end of the treatment after concentrating 3 L of the air exiting from the cuvette in a cartridge filled with 30 mg of Tenax and 30 mg of Carboxen (Gerstel, Mülheim an der Ruhr, Germany). A pump (Elite 5; A.P. Buck, Orlando, FL, USA) set at 200 ml min^−1^ rate was used to fill cartridges with the same volume of air without contamination from air that did not pass through the cuvette. All cartridges were stored at 4°C before being analysed through thermo-desorption followed by gas chromatograph-electro impact mass spectrometry (7890 GC – 5975 MSD 8 Agilent Tech, Santa Clara, CA, USA), as reported in Pollastri *et al.* (2018). Isoprene was identified by using the NIST 11.L 08 library spectral database and quantified with an isoprene gas standard (99.9%, Sigma-Aldrich) prepared in the laboratory.

During isoprene collection, a charcoal filter (Supelco, Bellafonte, USA) was placed ahead of the Li-Cor 6400 in order to remove all volatile organic compounds (including isoprene) from ambient air before reaching the gas exchange cuvette. Isoprene background was measured every day before starting the measurements by collecting 3 L of air the air exiting the empty cuvette.

### RNA sequencing

The first leaf was collected at the end of the treatment for RNA extraction and stored at - 80°C. RNA extraction was done with TRIzol^®^ Reagent (Ambion). RNA concentrations and quality were determined with NanoDrop spectrophotometer (Thermo Scientific, Wilmington, USA) and Agilent 2100 Bioanalyzer (Agilent, Santa Clara, CA). According to RNA quality and quantity, three out of four samples for each treatment were chosen for RNA sequencing and sent to the HuGeF sequencing service (http://www.hugef-torino.org, Human Genetics Foundation, Turin, Italy). A total of 15 paired-end libraries (2×75bp) were constructed using the TruSeq RNA library Prep Kit v2 (Illumina, San Diego, CA) with poly-A enrichment and sequenced on Illumina NextSeq 500. Raw data have been deposited in the Sequence Read Archive (https://www.ncbi.nlm.nih.gov/sra) with SRA accession SRP145569. Assessment of read quality metrics was carried out with FastQC software (http://www.bioinformatics.babraham.ac.uk/projects/fastqc/ version 0.11.3). Quality filtering, adapter cutting and trimming were carried out with Trimmomatic (version 0.33) (Bolger *et al.*, 2014), which can handle fastqc paired-end synchronization. After Illumina adapters clipping, the first 12 bases were trimmed due to sequencing biases (Hansen *et al*. 2010), leading and trailing low quality (below 3) or N bases and reads with low average quality (15) in a 4-bases scan were removed. Finally, reads less than 36 bases long after these steps were dropped.

Trinity software (version 2.0.6) was used for transcripts reconstruction. Contigs less than 200 bp and with coverage less than 5 were discarded (Haas *et al.* 2013). Transcripts redundancy was reduced with CD-HIT software (version 4.6.6), using a word size of 10 and 95% identity (Li and Godzik, 2006). Trinity software was able to assemble a total of 184,849 transcripts (Table S1). After removing redundancy, we obtained a total of 120,553 transcripts. The quality of our reconstructed transcriptome was tested mapping each paired-end library against it; the percentage of reads mapping back to transcriptome (RMBT) were between 78.66% and 89.87%, perfectly concordant with the expected percentage for RNA-seq experiment. In fact, because of the lack of a reference genome, the percentage of multimapping appears greater then single mapping (Table S2). Based on these results, the assembled transcriptome was considered reliable as reference for differential expression analysis.

### Identification of differentially expressed genes (DEGs), functional annotation and enrichment analysis

Reads from each of the 15 libraries were mapped against our reference transcriptome and quantified using RSEM (version 1.3.0) (Li and Dewey, 2011). The quantification was obtained as fragment per kb of exon per million fragments mapped (FPKM). In order to identify the differentially expressed genes, Trinity provided scripts based on the R package edgeR (R version 3.3.2; edgeR version 3.16.5) were used. Pairwise comparisons were made to highlight different expression in different conditions. Genes with a false discovery rate cutoff of 0.05 (5% FDR) were considered as differentially expressed.

To annotate the genes, blastx searches were performed against NCBI non-redundant database with an e-value cut-off of 1e^-3^. Blastx results, saved as xml files, were loaded into Blast2GO tool (version 4.1; database Germany, DE3, version b2g_jan17) (Conesa and Götz, 2008), and mapping, annotation and InterPro scanning were performed. To associate annotations obtained with Blast2GO to DEGs, the R package Annotation Tools (version 1.44.0) was used (Kuhn *et al.*, 2008). GO enrichment analysis of DEGs was performed on Blast2GO applying Fisher’s Exact Test with a FDR of 0.05. Pathways enrichment analysis of DEGs was carried out with KOBAS tool (v3.0) (Xie *et al.*, 2011) using *Oryza sativa* var. *japonica* as reference. Pathways were visualized with KEGG Mapper, a collection of tools for KEGG mapping (Kanehisa *et al.*, 2016).

### Quantification of metabolites

Soluble carbohydrates were identified and quantified by HPLC-RI analysis at the end of the treatment following the protocol of Tattini *et al.* (1996). Starch was quantified as reported in Chow and Landhäusser (2004) on the pellet resulting from ethanol extraction for the analysis of soluble carbohydrates. Glucose was quantified through peroxidase-glucose oxidase/o-dianisidine reagent (Sigma-Aldrich, Milano, Italy), reading the absorbance at 525 nm after the addition of sulfuric acid.

Hydrogen peroxide was measured spectrophotometrically at the end of the treatment after reaction with KI, according to a slightly modified method (Alexieva *et al.*, 2001). A modification of the Sedlak and Lindsay (1968) method was used for the glutathione (GSH) determination at the end of the treatment. Individual carotenoids were identified and quantified at the end of the treatment as reported in García-Plazaola and Becerril (1999). Phenylpropanoids were extracted and purified at the end of the treatment following the protocol of Tattini *et al.* (2004). Abscisic acid (ABA) was extracted and quantified at the end of the treatment using the protocol of López-Carbonell *et al.* (2009).

### Hydrogen peroxide (H_2_O_2_) localization and leaf ultrastructure by Transmission Electron Microscopy (TEM)

Hydrogen peroxide localization in leaves was estimated cytochemically via determination of cerium perhydroxyde upon reaction of cerium chloride (CeCl_3_) with endogenous H_2_O_2_, following the protocols of Bestwick *et al.* (1997) and Ranieri *et al.* (2003). At the end of the treatment, portion of approximately 0.15 mm^2^ were sampled in the center of the leaf blade and then infiltrated (under vacuum) with 5 mM CeCl_3_ in 50 mM 3-(N-morpholino)-propane sulfonic acid (pH 7.2). The CeCl_3_-treated and control leaf samples (without CeCl_3_-staining) were then fixed in 2.5% glutaraldehyde, in 0.2 M phosphate buffer (pH 7.2) for 1 h, and washed twice with the same buffer, prior of post-fixing with 2% osmium tetroxide in phosphate buffer (pH 7.2). Leaves were dehydrated in a graded ethanol series (30, 40, 50, 70, 90 and 100%), and gradually embedded in Spurr Resin (Sigma Aldrich). Ultrathin sections were obtained on an LKB IV ultramicrotome, mounted on Formvar coated copper grids, stained with UranylessEm Stain (Electron Microscopy Science) and lead citrate, and examined by using Philips CM12 transmission electron microscope (Philips, Eindhoven, The Netherlands) operating at 80 kV.

### Statistical analyses

Analysis of variance (ANOVA) was applied to test the effect of Na^+^ and P supply in *A. donax* plants. LSD post-hoc test was applied to assess significantly different means among treatments (P < 0.05 level).

## RESULTS

### Plant biometrics, gas exchange, chlorophyll fluorescence measurements and isoprene emission

At the end of the experiment, P concentration doubled in +P leaves, but did not decrease significantly in –P leaves, as compared to control (Table 1). In leaves of +Na plants, sodium (Na^+^) was two orders of magnitude higher than in control. When Na^+^ and P were both provided in excess (+NaP), an increase of Na^+^ and a slightly reduced accumulation of P, in comparison to leaves of +Na and +P plants, respectively, was observed.

**Table 1.** Plant biometrical traits (culm length and leaf number), relative water content (RWC), carbon, nitrogen (N), sodium (Na) and phosphorous (P) contents in leaves of *Arundo donax* plants in control conditions (C), without phosphorous supply (−P), with excess supply of phosphorous (+P) or sodium chloride (+Na), and with excess supply of both phosphorous and sodium chloride (+NaP). Data are means of 4 plants per treatment ± SE; different letters indicate statistical difference at P < 0.05 in the same column.

| | culm lenght<br>(cm) | leaf number<br>(n) | RWC<br>(%) | carbon<br>(%) | N<br>(%) | Na<br>($\mu\text{g g}^{-1}$ DW) | P<br>($\mu\text{g g}^{-1}$ DW) |
| --- | --- | --- | --- | --- | --- | --- | --- |
| C | 90.8 ± 2.0<sup>a</sup> | 16.0 ± 0.2<sup>ab</sup> | 89.6 ± 2.1<sup>a</sup> | 42.4 ± 0.3<sup>a</sup> | 3.4 ± 0.2<sup>a</sup> | 120.9 ± 23.1<sup>c</sup> | 2358.7 ± 168.4<sup>c</sup> |
| -P | 81.54 ± 2.9<sup>b</sup> | 15.2 ± 0.5<sup>b</sup> | 90.1 ± 0.6<sup>a</sup> | 42.4 ± 0.3<sup>a</sup> | 2.9 ± 0.2<sup>b</sup> | 103.5 ± 31.8<sup>c</sup> | 1766.7 ± 154.4<sup>c</sup> |
| +P | 91.3 ± 4.3<sup>a</sup> | 16.8 ± 0.5<sup>a</sup> | 91.5 ± 0.6<sup>a</sup> | 41.3 ± 0.6<sup>a</sup> | 3.7 ± 0.1<sup>a</sup> | 151.2 ± 16.3<sup>c</sup> | 5765.4 ± 403.2<sup>a</sup> |
| +Na | 62.6 ± 3.6<sup>c</sup> | 11.5 ± 0.3<sup>c</sup> | 83.0 ± 2.7<sup>b</sup> | 40.3 ± 0.5<sup>b</sup> | 3.8 ± 0.1<sup>a</sup> | 15098.3 ± 1128.1<sup>b</sup> | 2100.7 ± 109.0<sup>c</sup> |
| +NaP | 66.4 ± 2.5<sup>c</sup> | 10.7 ± 0.3<sup>c</sup> | 82.3 ± 1.1<sup>b</sup> | 39.5 ± 0.1<sup>b</sup> | 3.4 ± 0.1<sup>a</sup> | 17781.4 ± 1567.9<sup>a</sup> | 4501.3 ± 297.2<sup>b</sup> |

Excess supply of Na^+^ reduced culm length, number of leaves, leaf RWC, and leaf carbon content with respect to control (Table 1). P starvation reduced culm height, leaf number and nitrogen concentration, while P excess did not significantly affect any of the investigated parameters. However, in +NaP plants, culm length, number of leaves, leaf RWC and carbon content decreased to the same extent as in the +Na plants, with respect to control (Table 1).

Photosynthesis of *A. donax* decreased in –P, whereas it was similar to control in +P plants (Table 2). Photosynthesis was inhibited in +Na plants with respect to control, and the effect was even stronger in +NaP plants. In both +Na and +NaP plants, photosynthesis reduction was associated to reduced gs, Ci, and ETR, compared to control (Table 2). Isoprene emission from *A. donax* leaves was not affected by lack of P but was inhibited in +P plants, in comparison to control (Table 2). Isoprene emission was slight, but non statistically significant, stimulated by the +Na and +NaP treatments. (Table 2).

**Table 2.** Photosynthesis (A), O_2_ inhibition of photosynthesis (%), stomatal conductance (g_s_), internal CO_2_ concentrations (Ci) electron transport rate (ETR), isoprene emission of *Arundo donax* plants in control (C) conditions, without phosphorous supply (−P), with excess supply of phosphorous (+P) or sodium chloride (+Na), and with excess supply of both phosphorous and sodium chloride (+NaP). Data are means of 4 plants per treatment ± SE; different letters indicate statistical difference at P < 0.05 in the same column.

| | A<br>($\mu\text{mol m}^{-2} \text{s}^{-1}$) | Inhibition of<br>photosynthesis<br>(%) | g<sub>s</sub><br>($\text{mol m}^{-2} \text{s}^{-1}$) | Ci<br>(ppm) | ETR | Isoprene<br>($\text{nmol m}^{-2} \text{s}^{-1}$) |
| --- | --- | --- | --- | --- | --- | --- |
| C | 23.4 ± 1.1<sup>a</sup> | 29.9 ± 3.7<sup>b</sup> | 0.32 ± 0.03<sup>a</sup> | 239 ± 11<sup>a</sup> | 159 ± 5<sup>a</sup> | 17.9 ± 2.8<sup>a</sup> |
| -P | 18.2 ± 0.4<sup>b</sup> | 18.3 ± 2.8<sup>b</sup> | 0.24 ± 0.03<sup>a</sup> | 245 ± 16<sup>a</sup> | 140 ± 7<sup>a</sup> | 20.7 ± 4.4<sup>a</sup> |
| +P | 21.4 ± 1.2<sup>a</sup> | 26.2 ± 2.7<sup>b</sup> | 0.30 ± 0.05<sup>a</sup> | 250 ± 11<sup>a</sup> | 149 ± 6<sup>a</sup> | 12.0 ± 2.9<sup>b</sup> |
| +Na | 12.5 ± 2.0<sup>c</sup> | 48.3 ± 9.5<sup>a</sup> | 0.08 ± 0.03<sup>b</sup> | 167 ± 14<sup>b</sup> | 103 ± 19<sup>b</sup> | 25.9 ± 5.4<sup>a</sup> |
| +NaP | 4.0 ± 0.5<sup>d</sup> | 49.9 ± 12.2<sup>a</sup> | 0.03 ± 0.00<sup>c</sup> | 158 ± 21<sup>b</sup> | 71 ± 5<sup>b</sup> | 20.7 ± 3.3<sup>a</sup> |

### Analysis of differentially expressed genes (DEGs)

Exposure to high P concentration induced differences in the expression of a higher number of genes in *A. donax* leaves with respect to P starvation (Fig. 1, Table 3). The excess supply of P caused the differential expression of a similar number of up- and down-regulated genes, while the –P treatment mainly induced gene down-regulation. High concentration of Na^+^ had a higher impact on the total amount of DEGs (Fig. 1, Table 3), resulting in a higher extent of down-regulated genes with respect to high P treatment (Fig. 2). However, the number of DEGs increased 10-fold in +NaP treated plants (Fig. 1, Table 3) indicating, at molecular level, a higher response of *A. donax* to the combined (Na^+^ and P) than to the singularly applied treatments (Fig. 1, Fig. 2, Table 3). The complete list of DEGs is reported in Table S3.

**Table 3.** Number of differentially expressed genes (DEGs) at 5% FDR; (−P) low phosphorous, (+P) excess of phosphorous, (+Na) excess of sodium chloride, (+NaP) excess of both phosphorous and sodium chloride, (C) control condition.

|  | **-P vs C** | **+P vs C** | **+Na vs C** | **+NaP vs C** |
| --- | --- | --- | --- | --- |
| Up-regulated genes | 75 | 258 | 243 | 2103 |
| Down-regulated genes | 234 | 198 | 453 | 3076 |
| **Total number of DEGs** | **309** | **456** | **696** | **5179** |

**Figure 1.**
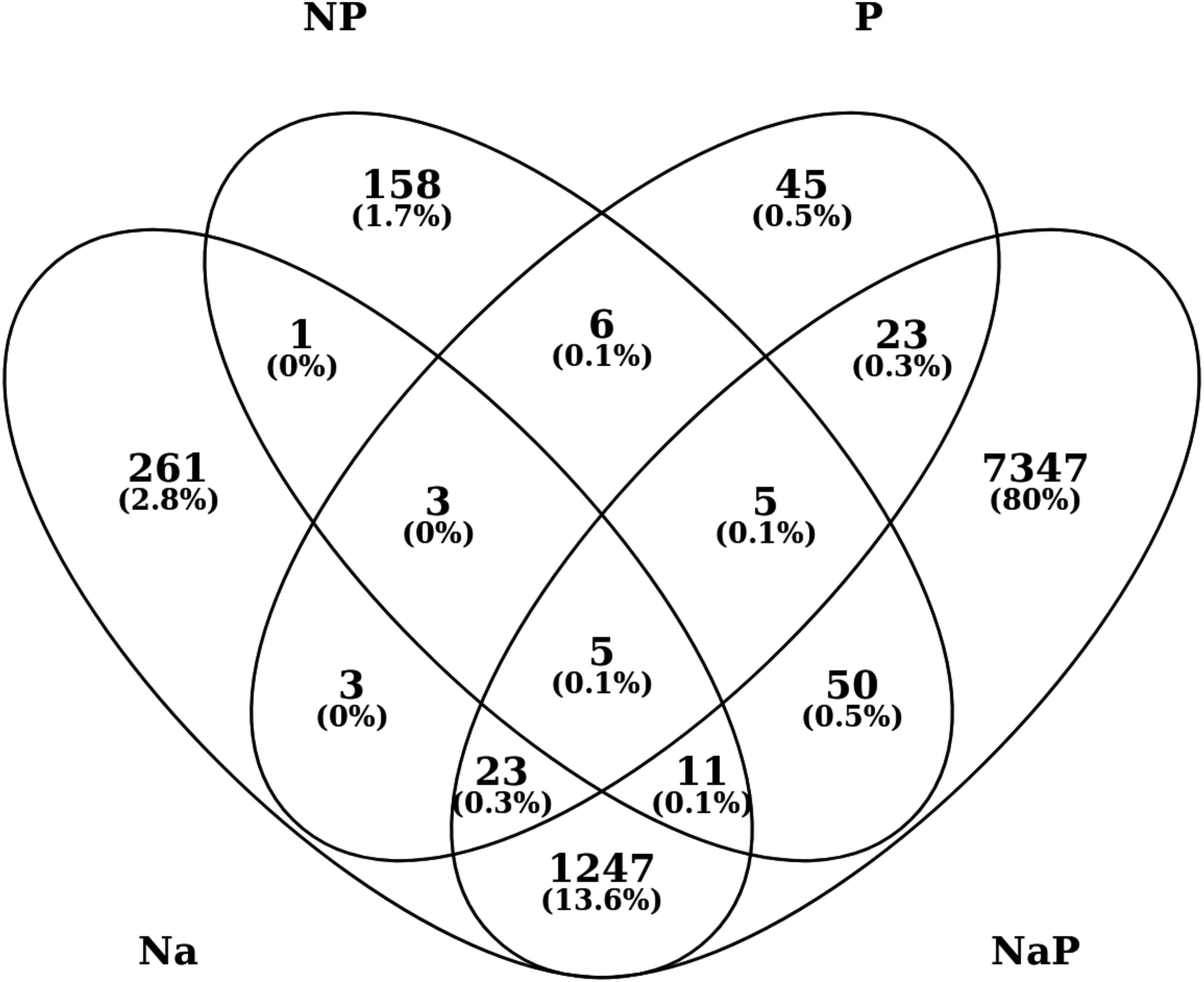
Venn diagram of differentially expressed genes (FDR<0.05) in *Arundo donax* plants without phosphorous supply (−P), with excess supply of phosphorous (+P) or sodium chloride (+Na), and with an excess supply of both phosphorous and sodium chloride (+NaP) with respect to control condition.

**Figure 2.**
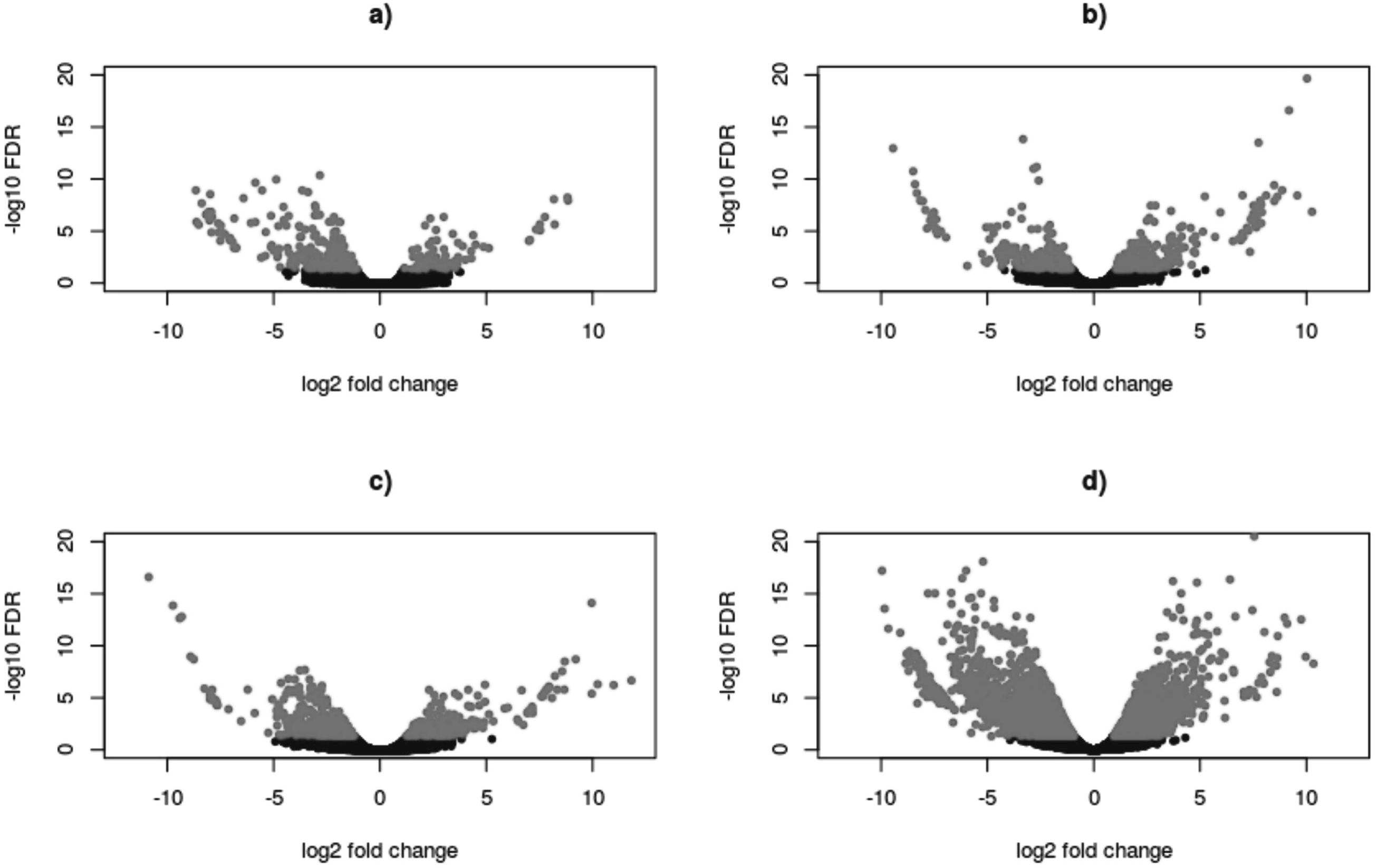
Volcano plots showing the entity of differentially expressed genes (DEGs) in *Arundo donax* plants without phosphorous supply (a: −P), with excess supply of phosphorous (b: +P) or sodium chloride (c: +Na), and with an excess supply of both phosphorous and sodium chloride (d: +NaP) with respect to control conditions. The log_2_-fold change (logFC) for each gene is plotted against log_10_-fold Fold Discovery Rate (logFDR). Significantly DEGs at 5% FDR are highlighted in grey.

In order to functionally inspect the overall DEGs and identify the major biological processes affected by the different supply of P and Na^+^, the transcriptome of *A. donax* leaves was annotated by mean of Gene Ontology (GO). More than half of the transcripts were annotated to at least one GO term (Fig. S1) and the first ten top-hit species found through the blastx search belonged to the *Poaceae* family (Fig. S2), indicating the reliability of the obtained GO annotation.

As a result of the GO category enrichment analysis (considering a p-value threshold of 0.05), only one functional category (‘*catalytic activities*’) was significantly over-represented in –P. Whereas, the over-represented categories were 33 (especially ‘*metabolic processes*’ and ‘*localization and transport*’) in +P, 38 (especially ‘*metabolic and biosynthetic processes*’ involving *‘protein binding’*, ‘*translocation and transportation’*, as well as ‘*catalytic activities’* and ‘*biological processes of the extracellular region’*) in +Na, and 139 (especially ‘*cellular, metabolic and biosynthetic processes of macromolecules and organic compounds*’ and ‘*binding activities*’, also involving the ‘*development of anatomical structure*’ and the ‘*organization of cellular (and intercellular) parts and organelles’*, as well as ‘*changes in the extracellular region*’) in +NaP plants. A complete overview of all the over-represented functional categories is shown in Table S4.

### Quantification of soluble carbohydrates and starch, photosynthetic pigments, abscisic acid (ABA), hydrogen peroxide (H_2_O_2_), glutathione (GSH) and caffeic acid derivative

Carbohydrate biosynthesis was impaired by different supply of P. However, a reduction of starch content was found in +P leaves, while in –P leaves the content of sucrose, fructose and non-structural carbohydrates was reduced compared to control (Table 4). DEG analysis showed that genes coding for ADP-glucose pyrophosphorylase, soluble acid invertases and a sucrose-phosphate synthase, involved in starch and sucrose metabolism pathway, were down-regulated in both treatments (Table S5). On the opposite, in +Na leaves the content of sucrose doubled compared to control, and +NaP treatment further increased the sucrose content and enhanced two-fold the contents of glucose, fructose, non-structural carbohydrates and starch, compared to control (Table 4). Moreover, pathway analysis on +NaP plants revealed an up-regulation of genes coding for enzymes involved in fructose and glucose synthesis (Table S5, Fig. S3B).

**Table 4.** Carbohydrates and starch content in leaves of *Arundo donax* plants in control conditions (C), without phosphorous supply (−P), with excess supply of phosphorous (+P) or sodium chloride (+Na), and with an excess supply of both phosphorous and sodium chloride (+NaP). Data are means of 4 plants per treatment ± SE; different letters indicate statistical difference at P < 0.05 in the same column.

|  | sucrose<br>(mg g<sup>-1</sup> DW) | glucose<br>(mg g<sup>-1</sup> DW) | fructose<br>(mg g<sup>-1</sup> DW) | non-structural<br>carbohydrates<br>(mg g<sup>-1</sup> DW) | starch<br>(mg g<sup>-1</sup> DW) |
| --- | --- | --- | --- | --- | --- |
| C | 4.4 ± 0.4<sup>c</sup> | 14.9 ± 0.5<sup>b</sup> | 67.8 ± 2.9<sup>b</sup> | 87.0 ± 3.6<sup>b</sup> | 5.2 ± 0.2<sup>b</sup> |
| -P | 3.3 ± 0.2<sup>c</sup> | 11.6 ± 0.5<sup>b</sup> | 45.1 ± 0.9<sup>c</sup> | 60.0 ± 14.5<sup>c</sup> | 4.8 ± 0.4<sup>bc</sup> |
| +P | 4.2 ± 0.3<sup>c</sup> | 14.2 ± 0.9<sup>b</sup> | 73.0 ± 4.6<sup>b</sup> | 91.4 ± 4.2<sup>b</sup> | 1.8 ± 0.5<sup>c</sup> |
| +Na | 11.4 ± 0.3<sup>b</sup> | 15.0 ± 0.8<sup>b</sup> | 71.4 ± 1.7<sup>b</sup> | 97.8 ± 2.7<sup>b</sup> | 4.3 ± 0.3<sup>b</sup> |
| +NaP | 14.1 ± 0.6<sup>a</sup> | 37.3 ± 2.7<sup>a</sup> | 166.0 ± 10.6<sup>a</sup> | 217.4 ± 13.1<sup>a</sup> | 7.1 ± 0.2<sup>a</sup> |

In +Na and +NaP leaves, leaf ABA content increased two-fold with respect to control (Fig. 3C). In these same leaves, molecular analysis showed a down-regulation of the gene coding for the ABA 8-hydroxylase 3, a key enzyme in ABA catabolism (Table S3). However, three ABA stress-ripening coding genes, involved in response to abiotic stress, were induced in +NaP plants (Table S3).

**Figure 3.**
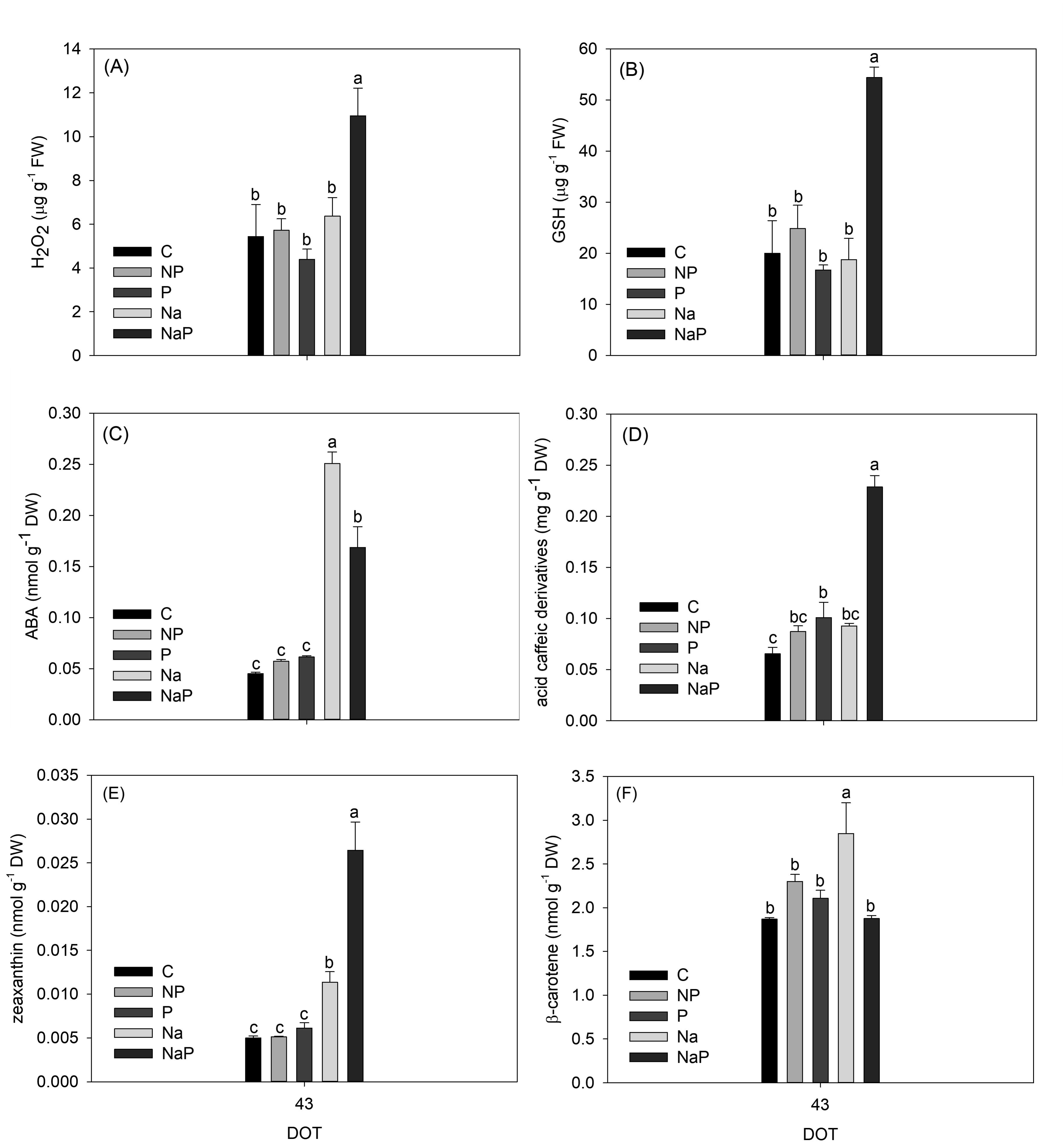
Hydrogen peroxide (H_2_O_2_) (A), glutathione (GSH) (B), abscisic acid (ABA) (C), caffeic acid derivative (D), zeaxanthin (E) and β-carotene (F) contents of *A. donax* plants in control (C) conditions, without phosphorous supply (−P), with excess supply of phosphorous (+P) or of sodium chloride (+Na), and with an excess supply of both phosphorous and sodium chloride (+NaP). Data are means of 4 plants per treatment ± SE; different letters indicate statistical difference at P < 0.05.

Hydrogen peroxide (H_2_O_2_) and glutathione (GSH) highly accumulated in +NaP plants with respect to all the other treatments (Fig. 3A, B). Consistent with these observations, genes involved in the glutathione metabolism were more up-regulated in +NaP leaves than in +P leaves (Fig. S3E, Fig. S3F).

The content of flavonoids was significantly enhanced in +NaP leaves, while the other treatments caused only a moderate increase of these secondary metabolites, with respect to control (Fig. 3). The pathway of flavonoids biosynthesis was significantly perturbed in +NaP plants. Indeed, genes like flavonol synthase, trans-cinnamate 4-monooxygenase, flavonoid 3′-monooxygenase and chalcone synthase were up-regulated in +NaP plants with respect to control (Fig. S3C, Fig. S3D).

Zeaxanthin and β-carotene were enhanced in +Na leaves with respect to control, whereas zeaxanthin and acid caffeic derivatives were further stimulated in +NaP plants (Fig. 3C, D, E, F). Although there was no differential regulation in genes involved in zeaxanthin and β-carotene synthesis in +Na leaves, a down-regulation of lycopene β-cyclase and phytoene synthase, genes responsible for β-carotene synthesis was measured in +NaP plants (Table S3).

### Transmission electron microscopy images of leaves ultrastructure

The ultrastructure of *A. donax* control leaves highlighted a peripheral location of organelles, and a large vacuole in the center of the cells (Fig. 4). Cytoplasmic organelles (nucleus, mitochondria, vacuole, endoplasmic reticula, Golgi apparatus) showed typical structure and distribution, and chloroplasts had distinct granal and stromal thylakoid arrangement and a well-defined stroma matrix where few and little starch grains were present (Fig. 4B).

**Figure 4.**
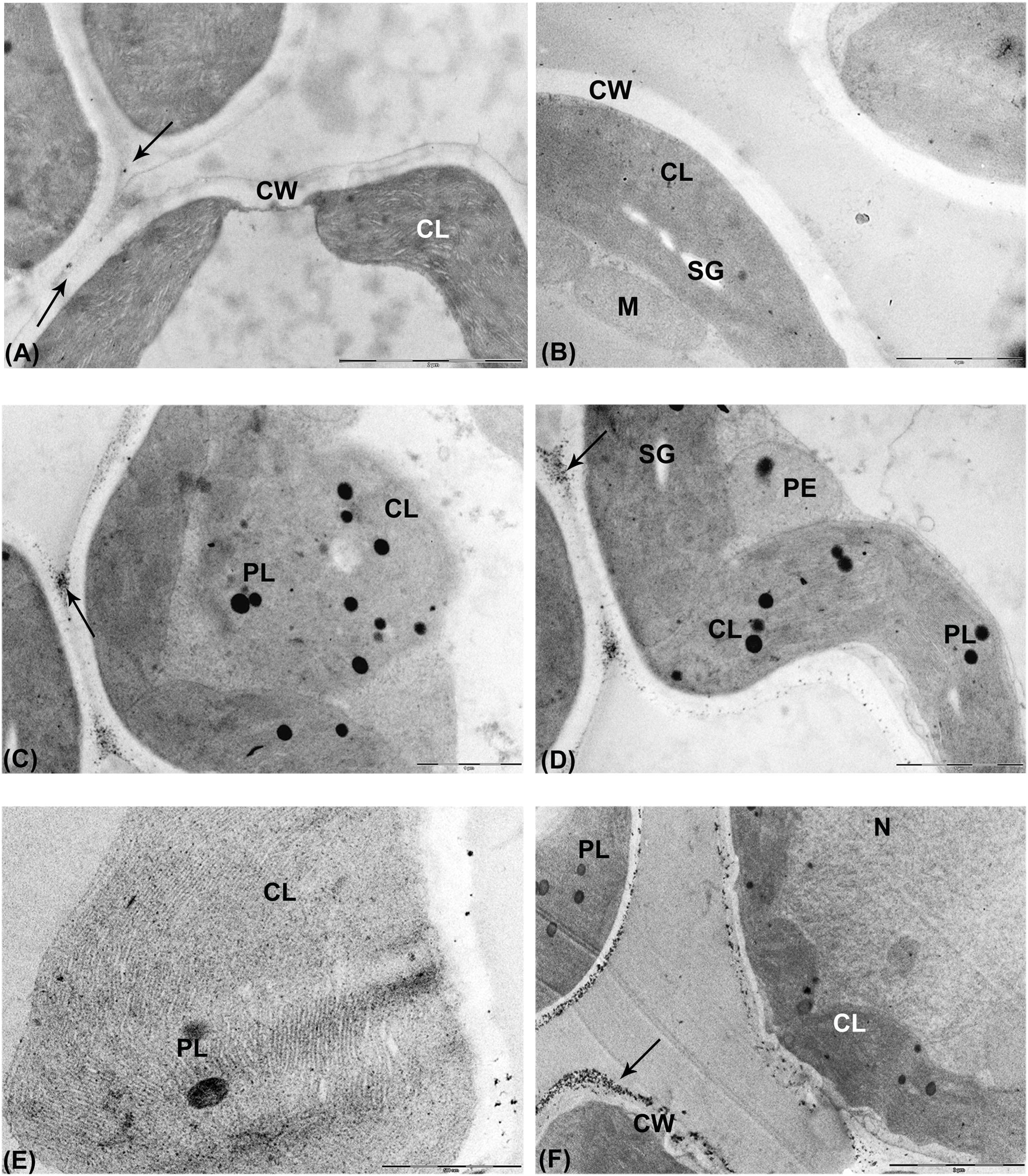
Micrographs of leaf ultrastructure of *A. donax* in control conditions with CeCl_3_ (A) and without CeCl_3_ (B), with excess supply of phosphorous (C, D) and without phosphorous (E, F) supply. Legend: CL: chloroplast; CW: cell wall; L: lipid body; M: mitochondrion; N: nucleus; PE: peroxisome; PL: plastoglobule; SG: starch grain; V: vacuole. Black arrows refer to electron-dense deposits of CeCl_3_, indicative of the presence of H_2_O_2_. A, B, C, D: bar 1 µm; E: bar 100 nm; F: bar 2 µm.

In +P plants, chloroplasts displayed very little or no starch grains (Fig. 4C), confirming the decrease of starch also reported in Table 3. These cells showed more and bigger plastoglobules than those of control plants (Fig. 4C). In addition, the envelope membrane of few +P chloroplasts appeared damaged, with thylakoids not clearly recognizable (Fig. 4C). Some +P cells also had wavy plasma membrane, large peroxisomes (Fig. 4D), and electron dense cerium perhydroxide precipitates in the cell walls after treatment with CeCl_3_, thus indicating the onset of ROS accumulation and stressful conditions (Fig. 4C; 4D).

Chloroplasts of –P leaves were characterized by an extensive system of grana and stroma lamellae (also reported by Hall et al., 1972), filling the stroma, that also contained a moderate number of plastoglobules (Fig. 4E). The nucleus of –P cells showed poorly condensed chromatin. Moreover, deposition of CeCl_3_ was found in cell walls (Fig. 4F) and in bundle sheath cells.

In leaves of +Na plants, the shape of the cells changed from elliptical to wrinkle elongated, and cell walls appeared curled (indicated by arrows in Fig. 5A). Strong local H_2_O_2_ accumulation in the cell walls (indicated by the black arrow in Fig. 5B) and large cytoplasmic lipid bodies (Fig. 5B) were detectable in some +Na cells. Moreover, some mesophyll cells were destroyed, and cytoplasmic organelles were no longer recognizable except for swollen or disintegrated chloroplast (Fig. 5C). However, chloroplasts of +Na cells that were still visible showed a wavy outline (indicated by the white arrow in Fig. 5B), significant loss of clear stromal matrix, with swelling and curling thylakoids, and an increased number of plastoglobules (Fig. 5A, 5B). In addition, many peroxisomes with scarce electron dense deposits of CeCl_3_ were observed (data not shown).

**Figure 5.**
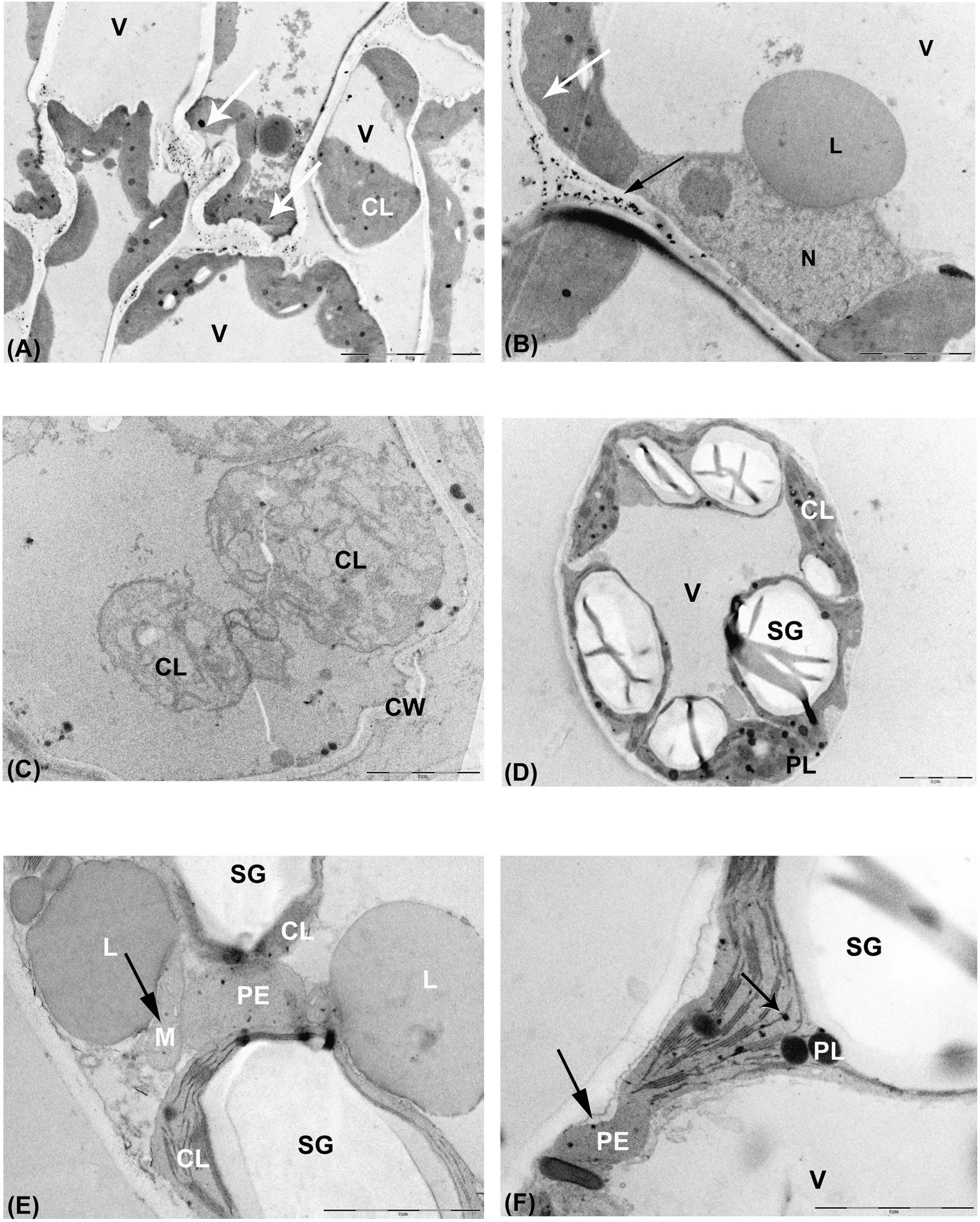
Micrographs of leaf ultrastructure of *A. donax* with excess supply of sodium chloride (A, B, C), with both excess supply of phosphorous and sodium chloride (D, E, F). Legend: CL: chloroplast; CW: cell wall; L: lipid body; M: mitochondrion; N: nucleus; PE: peroxisome; PL: plastoglobule; SG: starch grain; V: vacuole. White arrows refer to wavy structure; black arrows refer to electron-dense deposits of CeCl_3_, indicative of the presence of H_2_O_2_. A: bar 5 µm; B, C, D: bar 2 µm; E, F: bar 1 µm.

Mesophyll cells of +NaP plants contained chloroplasts with numerous and large plastoglobules and very large starch granules (Fig. 5D), matching the reported increase of starch (Table 3) and carotenoid (i.e., zeaxanthin) content of these leaves (Fig. 2E). Large lipid bodies were also present in the cytoplasm of these cells, and CeCl_3_ deposits were observed in chloroplasts, peroxisomes and mitochondria (Figs. 5E; 5F).

## Discussion

### Performance of *Arundo donax* grown under high- or low-concentration of phosphorus (P)

A two-fold increase of P in the leaves of *A. donax* approached toxic levels, as confirmed by early symptoms of alteration of cell ultrastructure and the presence of peroxisomes (Fig. 4D), indicating starting oxidation processes. However, high P concentrations did not hamper *A. donax* growth and photosynthesis (Table 1). Tolerance of photosynthesis to high P concentration could be the result of the tight regulation of P homeostasis within the cytoplasm, due to the activation of mechanisms that transport and store the excess of P into the vacuoles (Mimura *et al.*, 1990). However, excess of P strongly decreased starch accumulation in leaves (Table 4), as confirmed by histological observations (Fig. 4C, D). Our transcriptomics results indicate that the inhibition of starch metabolism in *A. donax* exposed to +P was mainly due to the transcriptional repression of the ADP-glucose pyrophosphorylase, rather than by enhanced translocation of triosophosphates, that reduces the availability of these substrates for starch synthesis in the chloroplasts (Pozueta-Romero *et al*., 1991; Heldt *et al*., 1991). Moreover, in +P plants there was a strong induction of few transcripts coding for cytosolic fructose-1,6-bisphosphatase an enzyme that, by catalyzing the first irreversible reaction that turn fructose-1,6-bisphosphate into fructose-6-phosphate and inorganic phosphate (Ladror *et al*., 1990), plays an important regulatory role in carbohydrates biosynthesis and metabolism (Daie, 1993).

Consistently with previous results (Fares *et al*., 2008) and a recent meta-analysis (Fernández-Martínez *et al*., 2017), leaves of +P plants emitted less isoprene than control and –P plants. Although isoprene production is a highly ATP demanding process (Loreto and Sharkey 1990), exposure to high P concentration may prompt a competition between mitochondrial respiration and the methylerythritol 4-phosphate (MEP) pathway, in turn limiting isoprene biosynthesis (Loreto *et al*., 2007). In particular, phosphoenolpyruvate (PEP) is a substrate for both isoprene biosynthesis and mitochondrial respiration. Mitochondrial respiration was likely stimulated in plants grown at high P concentration (Fares *et al*. 2008). Indeed, we observed an increased transcription of genes involved in energy requiring processes of protein production and export (Fig. S3). Therefore, our results seem to indicate that incorporation of P into PEP, principally serving the respiratory metabolism, made it less available for isoprene production.

*A. donax* was sensitive to P deficiency. Although 43 days of P starvation did not significantly decrease the leaf P concentration, in −P leaves the expression of numerous genes was down-regulated (Müller *et al*., 2007; Hernández *et al*., 2007) and the ultrastructure of leaf cells was altered. Further results confirmed that, under reduced P availability, *A. donax* reduces photosynthesis, grows shorter, and produces a lower number of leaves (with reduced N content) than plants grown under normal P availability. Sensitivity to low P availability may affect the capacity of *A. donax* to colonize new habitats (Wassen *et al*., 2005), and limits *A. donax* use for biomass production in poorly fertile soils.

### Different response of *A. donax* to Na^+^ stress, and to synergistic action of high Na^+^ and P

Accumulation of Na^+^ in leaves affected stomatal conductance by increasing diffusive limitations of photosynthesis (the acquisition of CO_2_ to be assimilated), as further confirmed by low values of Ci. This response to high Na^+^ concentrations widely occurs across plant species (Delfine *et al*., 1999; Centritto *et al*., 2003). Stomata closure was likely triggered by increased synthesis of ABA upon salinity stress (Wilkinson and Davies, 2002; Seiler *et al*., 2011). In leaves of Na^+^-stressed *A. donax*, photosynthesis and ETR were strongly reduced, whereas zeaxanthin and β-carotene were largely synthesized. This suggests the onset of coordinated photochemical processes to inhibit the accumulation of reactive oxygen species (ROS). Salinity stress also stimulated the biosynthesis of sucrose in *A. donax* leaves (Table 4), as also confirmed by the significant over-representation of GO categories related to ‘*carbohydrate metabolic process*’ (Table S4). Beside exerting a signaling role (Park *et al*., 2016), sucrose was likely able to balance the drop in osmotic potential as leaf RWC decreases during progressive exposure to salinity stress (Table 1). However, changes observed to the leaf ultrastructure of Na^+^-stressed plants, where only some mesophyll cells and chloroplast resulted completely destroyed (Fig. 5), confirmed that *A. donax* is moderately sensitive to high Na^+^ concentration in leaves. Indeed, *A. donax* possesses more glycophytic than halophytic features (Nackley and Kim, 2015) and tolerates salinity through mechanisms that may prevent ROS formation despite accumulation of Na^+^ in the leaves (Mumm and Tester, 2008).

Salinity impaired photosynthesis but increased (although not significantly) isoprene emission from *A. donax* leaves. Isoprene is synthesized from carbon assimilated through photosynthesis (Delwiche and Sharkey, 1993), but its emission may be also sustained by extra-chloroplastic carbon sources when photosynthesis is limited under (abiotic) stress (Brilli *et al.*, 2007; Fortunati *et al*., 2008). Overall, the simultaneous increases in the biosynthesis of isoprene and carotenoids may imply activation of the 2-C-methyl-D-erythritol 4-phosphate (MEP) pathway in +NaP-stressed *A. donax* plants to enhance protection against stressful condition (Loreto *et al*., 2014; Marino *et al*., 2017).

An additive effect of simultaneous supply of high Na+ and P concentrations was clearly highlighted by a 10-fold increase in the number of both up- and down-regulated genes in leaves of +NaP *A. donax* plants. Some of the most representative transcription factors already identified in *A. donax* under drought (Fu *et al*., 2016) were also regulated under Na^+^ and P stress. Among them, NAC was strongly induced in +Na, +P and +NaP, whereas WRKY 50, 53 and 41 were down-regulated only in +NaP plants. NAC and WRKY genes family are known to mediate water-(Hadiarto and Tran, 2011) and Na^+^-stress responses, as well as ABA signaling pathway in plants (Jiang *et al*., 2017). Genes coding for stress-associated proteins (SAPs) are important regulators of multiple abiotic stress tolerance (Giri *et al*., 2013) and found to be induced in water-stressed *A. donax* plants (Evangelistella *et al*., 2017). However, only two SAPs were down-regulated in +Na and +NaP plants. Despite inducing a higher expression of genes involved in abiotic stress tolerance (e.g., NAC, WRKY and SAP genes), high P concentration exacerbated the reduction of photosynthesis in Na^+^-stressed *A. donax* plants, as also indicated by the over-representation of many GO categories related to ‘*cellular metabolic process*’ in +NaP plants (Table S4). Photosynthesis could have been limited by altered sugar metabolism, as the amount of non-structural carbohydrates, fructose, glucose and starch increased two-fold in +NaP plants. It is suggested that combined supply of Na^+^ and P strongly reduced the turnover of carbohydrates, which may have favored the formation of large starch grains in the chloroplasts (Fig. 5D, E, F). Our results show that increase of starch biosynthesis in +NaP plants was related (as in +P plants) to the induction of the ADP-glucose pyrophosphorylase, whereas translocation of triosephosphates was not significantly affected. However, in +NaP plants photosynthesis was stimulated under low O_2_ conditions, indicating that feedback inhibition of photosynthesis, typically induced by carbohydrates accumulation (Sharkey, 1990; Xu *et al*., 2015) did not occur. The accumulation of carbohydrates induced by P supply in Na^+^-stressed plants may serve protective purposes, in enhancing osmotic capacity to assimilate water (Lambers *et al*., 2008) as confirmed by the over-representation of GO categories regarding ‘*organic substance metabolic process*’ in +NaP plants (Table S4). Carbohydrate accumulation may also help prevent damage to the cell structures (Yang and Guo, 2017). Indeed, the +NaP treatment induced a SNF1-related protein kinase coding gene (SnRK2), which was also found to be responsive to both ionic and non-ionic osmotic stressful conditions (Fu *et al*., 2016; Virlouvet and Fromm, 2015). Genes coding for dehydrins (DHNs) proteins, which play cellular protection in abiotic stress tolerance (Gao and Lan, 2016; Verma *et al*., 2017), were also up-regulated in leaves of +NaP plants. Remarkably we also observed that, while Early Responsive to Dehydration (ERD4) were induced as expected in *Poaceae* (see Fu *et al.*, 2016 for similar finding in drought-stress conditions), two ERD6 genes coding for carbohydrate transporters were down-regulated, consistent with carbohydrates accumulation shown in leaves of +NaP plants.

Interaction between high concentrations of Na^+^ and P did not significantly affect isoprene emission, which was once again uncoupled from photosynthesis in stressed leaves (Brilli *et al*., 2007; Vickers *et al*., 2009; Marino *et al*., 2017). The small reduction of isoprene emission with respect to +P leaves could be associated to the very large negative effect of the combined treatment (+NaP) on photosynthesis, and to the consequently reduced photosynthetic substrate entering the MEP pathway. Transcriptomics show up-regulation of ABA biosynthesis and down-regulation of β-carotene (both made by MEP) in +NaP compared to +Na leaves. This suggests a rearrangement of the flux of carbon into the MEP pathway towards hormones controlling stomata movement and away from antioxidants such as isoprene and carotenoids. Moreover, competition with starch for PEP could also limit isoprene synthesis in +NaP as well as in +P leaves (see above). However, starch was not a limiting factor in +NaP leaves as it was in +P leaves. The accumulation of carbohydrates in +NaP leaves was also highlighted by the over-expression of GO categories related to ‘*organic substance metabolic process*’ (Table S4), suggesting that a glucose 6-phosphate shunt might have been activated to increase the availability of precursors for the MEP pathway (Sharkey and Weise, 2017), despite the low flux of carbon fixed by photosynthesis.

In +NaP leaves, a significant increase of both H_2_O_2_ and glutathione (GSH) contents was observed, and further confirmed in our transcriptome analysis by over-representation of GO categories ‘*cellular metabolic process’ and ‘organic substance metabolic process*’ (Table S4), indicating enhanced ROS formation and activation of the anti-oxidant metabolism. Moreover, enhanced biogenesis of peroxisomes in +NaP leaf cells (Fig. 5E, F), most likely indicates a general increase of oxidative stress conditions (Lopez-Huertas *et al*., 2000). Peroxisomes contain antioxidants enzymes able to metabolize ROS and to enhance tolerance to a wide range of stresses (Nyathi and Baker, 2006). High synthesis of GSH possibly prevents the increase of H_2_O_2_ to reach toxic level while allowing this compound to exert signaling functions (Mittler, 2002; Baxter *et al*., 2014) that may further enhance stress response (Knight and Knight, 2001). However, this was clearly insufficient to protect photosynthesis in +NaP leaves.

## Conclusions

We showed that *A. donax* can be cultivated in marginal soils affected by eutrophication (under high P supply), where it can exert positive functions (e.g. for phytoremediation) despite allocating less carbon to defensive secondary metabolites (isoprene). Moreover, our results highlight that *A. donax* is sensitive to P deficiency and to Na^+^ excess, and this sensitivity is further enhanced by the combination of high Na^+^ and high P. However, the supply of high P concentrations further stimulates, at molecular and biochemical level, responses that favour stress tolerance in salt-stressed *A. donax*. Therefore, although the productivity of *A. donax* may be largely impaired, the plant adapts and survives in unfavourable soils rich in both P and Na^+^.

## Acknowledgements

The study was funded by research project “CROPSTRESS - System performance of non-food crops to drought stress: development of a plant ideotype”, by the SIR2014 program of the Italian Ministry of University and Research (RBSI14VV35), Claudia Cocozza is the scientific leader of the project. We thank for providing the instrumentation needed for isoprene Dr. Marco Michelozzi, Laboratory for the Analysis and Research in Environmental Chemistry (ARCA), and for Na and P determination Prof. Cristina Gonnelli.

## Author contributions

CC and BF designed the research, analyzed the data and wrote the article with contributions of all the authors; PiS, CC, BF performed research; ML, RS, BC, GC, PoS, MM provided technical assistance; CM, AG, TR, LF supervised and complemented the writing.

